# A new role of the Mus81 nuclease for replication completion after fork restart

**DOI:** 10.1101/785501

**Authors:** Benjamin Pardo, María Moriel-Carretero, Thibaud Vicat, Andrés Aguilera, Philippe Pasero

**Affiliations:** Institut de Génétique Humaine, CNRS / Université de Montpellier, Montpellier, France; Centro Andaluz de Biología Molecular y Medicina Regenerativa CABIMER, Universidad de Sevilla-CSIC-Universidad Pablo de Olavide, Seville, Spain; Centre de Recherche en Biologie cellulaire de Montpellier, CNRS / Université de Montpellier, Montpellier, France

**Keywords:** fork restart, Mus81, recombination, replication, termination, BIR

## Abstract

Impediments to DNA replication threaten genome stability. The homologous recombination (HR) pathway is involved in the restart of blocked replication forks. Here, we used a new method to study at the molecular level the restart of replication in response to DNA topoisomerase I poisoning by camptothecin (CPT). We show that HR-mediated restart at the global genomic level occurs by a BIR-like mechanism that requires Rad52, Rad51 and Pol32. The Mus81 endonuclease, previously proposed to cleave blocked forks, is not required for replication restart in S phase but appears to be essential to resolve fork-associated recombination intermediates in G_2_/M as a step necessary to complete replication. We confirmed our results using an independent system that allowed us to conclude that this mechanism is independent of the accumulation of DNA supercoiling and DNA-protein crosslinks normally caused by CPT. Thus, we describe here a specific function for Mus81 in the processing of HR-restarted forks required to complete DNA replication.

## INTRODUCTION

Chromosome duplication during the S phase is a crucial step of cell division. DNA replication in eukaryotes is initiated from multiple origins distributed on all chromosomes. Replication machineries progress along chromosomes arms by coupling the unwinding of the DNA double helix to the synthesis of DNA complementary to the template strands until merging with another incoming fork at termination regions. The progression of replication forks can be hindered by obstacles in the DNA template such as DNA lesions. Genome stability is particularly at risk when damaged DNA molecules are replicated. Failure in DNA damage repair can lead to the terminal arrest or breakage of replication forks, and ultimately to the distribution of under-replicated and / or broken chromosomes to the daughter cells after mitotic division. When a fork becomes dysfunctional, the completion of replication could be ensured by a converging functional fork, a process that can be favored by the firing of nearby dormant origins (Brambati et al., 2018, Yekezare et al., 2013). Alternatively, dysfunctional replication forks could also be restarted, a mechanism that requires the homologous recombination (HR) pathway (Ait Saada et al., 2018, Lambert et al., 2010, Mayle et al., 2015).

HR is activated during the S phase of the cell cycle (Aylon et al., 2004, Huertas et al., 2008), when sister-chromatids become available as repair templates (Gonzalez-Barrera et al., 2003, Kadyk & Hartwell, 1992). This has been extensively studied in the context of DNA double-strand break (DSB) repair (Pardo et al., 2009) but HR is also involved the repair of DNA lesions within single-strand DNA (ssDNA) gaps left behind the replication fork by the damage avoidance pathway (Branzei & Foiani, 2010). Classically, HR is characterized by the succession of three steps: 1) resection of the 5’-ended DNA strand at DSB ends or in ssDNA gaps, followed by 2) strand invasion of a homologous DNA duplex and priming of DNA synthesis through the formation of a displacement loop (D-loop), and 3) resolution of recombination intermediates (Paques & Haber, 1999). During HR, Rad52 and Rad51 are fundamental to coat the ssDNA generated by resection and carry out the strand invasion and exchange reactions (Krogh & Symington, 2004). At dysfunctional forks, Rad51 loading by Rad52 also regulates nascent DNA strands degradation by exonucleases (Ait Saada et al., 2017, Schlacher et al., 2011, Vallerga et al., 2015). This protection mechanism is thought to occur upon the displacement and reannealing of the nascent strands together, making replication forks to reverse (Meng & Zhao, 2017, Teixeira-Silva et al., 2017, Zellweger et al., 2015).

Recombination-mediated replication restart has been mainly studied in the yeast cellular model using site-specific tools to break or block replication forks (Cortes-Ledesma & Aguilera, 2006, Gonzalez-Barrera et al., 2003, Lambert et al., 2010, Mayle et al., 2015, Mohebi et al., 2015, Nielsen et al., 2009, Roseaulin et al., 2008). Replication restart in *Saccharomyces cerevisiae* has been recently investigated using a Flp-nick system, which produces a site-specific DNA nick that is converted into a one-ended DSB upon the passage of a replication fork (Jakobsen et al., 2019b, Mayle et al., 2015, Nielsen et al., 2009). Recombination-mediated restart seems to occur by break-induced replication (BIR), a HR pathway mainly characterized outside of S phase (Anand et al., 2013, Kramara et al., 2018). BIR is favored when the DSB only displays one end that is homologous to the repair template. In that case, DNA synthesis primed within the D-loop can proceed to the chromosome end by a conservative mechanism (Donnianni & Symington, 2013, Malkova et al., 2005, Morrow et al., 1997, Saini et al., 2013). In particular, the non-essential DNA polymerase δ subunit Pol32 and the Pif1 DNA helicase are required to promote the migration of the D-loop (Lydeard et al., 2007, Saini et al., 2013, Wilson et al., 2013). Lastly, BIR synthesis is highly mutagenic (Deem et al., 2011, Mayle et al., 2015) and subjected to template switching, leading to complex chromosomal rearrangements(Anand et al., 2014, Pardo & Aguilera, 2012, Ruiz et al., 2009, Smith et al., 2007). Terminally-blocked replication forks at the *RTS1* barrier in *Schizosaccharomyces pombe* are restarted without DSB formation by a Rad52- and Rad51-dependent mechanism, requiring both their fork protection and recombination functions (Ahn et al., 2005, Ait Saada et al., 2017, Lambert et al., 2010, Lambert et al., 2005, Nguyen et al., 2015, Teixeira-Silva et al., 2017). As for BIR in *S. cerevisiae*, replication restart at *RTS1* in *S. pombe* requires the DNA polymerase δ and the Pfh1 DNA helicase, is highly mutagenic and prone to template switching (Iraqui et al., 2012, Jalan et al., 2019, Miyabe et al., 2015, Nguyen et al., 2015).

Whereas numerous factors have been identified to participate in the first two steps of HR-mediated restart of DNA replication, namely 1) DNA resection and 2) strand protection/invasion, it remains unclear which factors participate to the resolution of the recombination intermediates in this context. The structure-selective endonuclease (SSE) Mus81 has been proposed to cleave either blocked or reversed forks to promote repair by HR (Hanada et al., 2007, Pepe & West, 2014, Regairaz et al., 2011). However, Mus81 is not required for replication restart at *RTS1* in fission yeast, nor for the repair of a replication-born DSB in budding yeast (Lambert et al., 2010, Roseaulin et al., 2008). Mus81 is nevertheless involved in the processing of these recombination events, as they accumulate in its absence (Lambert et al., 2010, Roseaulin et al., 2008), resulting in a decreased amount of final repair products (Munoz-Galvan et al., 2012, Roseaulin et al., 2008). By using the Flp-nick system, it has been proposed that Mus81 limits replication restart by Pol32-dependent BIR in S phase by processing the migrating D-loop (Mayle et al., 2015). Mus81 catalytic activity is normally very low in S phase, and only increases at the G_2_/M transition, when the Mms4 regulating subunit of the complex is hyper-phosphorylated by multiple kinases (Gallo-Fernandez et al., 2012, Matos et al., 2011, Matos et al., 2013, Princz et al., 2017). Hence, the role of Mus81 in replication restart by HR in S phase appears contradictory to the regulation of its activity. We took advantage of our study to clarify the role of Mus81 in recombination-mediated restart of DNA replication.

Finally, it remains to be determined if replication restart studied at locus-specific barriers occurs in the same way at the genome-wide level in response to natural replication impediments. The DNA topoisomerase 1 (Top1) normally introduces a transient nick to relax supercoiled DNA during transcription and replication (Pommier et al., 2016). During this reaction, Top1 remains covalently attached to the 3’ end of the break, forming a “cleavage complex” (Top1cc), before the relaxation of DNA and religation of the break. Cells are constantly challenged with blocked Top1ccs, which is a major driver of mutagenesis in highly transcribed genes (Lippert et al., 2011, Takahashi et al., 2011). Furthermore, cells devoid of both Tdp1 and Wss1, two factors involved in the removal of Top1ccs, have a severe Top1-dependent growth defect (Balakirev et al., 2015, Stingele et al., 2014), which indicates that Top1ccs are natural threats to cell survival. Top1ccs can also be stabilized by the drug camptothecin (CPT) (Pommier et al., 2010). CPT is thought to cause the formation a one-ended DSB, the typical substrate for BIR, upon the collision of the replication fork with the nick in the Top1cc, thus inducing a replication stress (Hsiang et al., 1989, Nitiss & Wang, 1988, Strumberg et al., 2000). However, CPT-induced DSBs have not been observed in yeast cells (Ray Chaudhuri et al., 2012, Redon et al., 2003). More recently, it has been shown that CPT treatment of yeast cells induces fork reversal (Menin et al., 2018, Ray Chaudhuri et al., 2012), as a possible consequence of accumulation of positive supercoils ahead of replication forks due to Top1 inhibition (Koster et al., 2007). Hence, further investigation is required to clarify the consequences of Top1 poisoning by CPT on replication forks.

Our data reveal that HR-mediated replication restart at Top1ccs likely occurs by Rad51- and Pol32-dependent BIR-like mechanism. Thanks to a method to increase CPT entry into yeast cells, we show for the first time that replication restart in response to Top1 poisoning requires Rad52 and Rad51 but not Mus81. Mus81 is rather required in G_2_/M to promote the termination of restarted forks. We confirmed our results in CPT-treated cells using an independent system in which a specific mutation of the Rad3 DNA helicase generates a similar replication stress. This allowed us to demonstrate that the mechanism we have characterized upon Top1 poisoning is independent of the accumulation of supercoiled DNA and DNA-protein crosslinks. The processing of HR-restarted forks in order to complete replication appears therefore as a general function of the Mus81 endonuclease.

## RESULTS

### Top1 poisoning by CPT and the *rad3-102* allele exert a genetically similar replication stress

In this study, we have chosen to study the dynamics of DNA replication challenged by CPT treatment, which exacerbates at the genome-wide level the presence of trapped Top1 on the DNA template. Resistance to CPT-induced DNA damage absolutely requires the homologous recombination (HR) machinery, as null mutations of *RAD52, MRE11*, *RAD50* and *XRS2* lead to the highest sensitivities to low doses of CPT compared to wild type cells (**Figure 1A**) (Liu et al., 2002, Vance & Wilson, 2002). *rad51*Δ mutants are also highly sensitive to low CPT doses but to a lesser extent than its upstream regulator Rad52 (**Figure EV1A**) (Mayle et al., 2015, Vance & Wilson, 2002). Interestingly, Rad52 and the members of the MRX complex (Mre11-Rad50-Xrs2), but not Rad51, are essential for the survival of cells bearing the *rad3-102* mutation: this mutation impairs the nucleotide excision repair (NER) pathway by increasing the binding of the TFIIH complex to a single-strand DNA gap intermediate, which prevents its subsequent filling (Herrera-Moyano et al., 2014, Moriel-Carretero & Aguilera, 2010a). Additionally, it has been described that the elevated CPT sensitivity of a nuclease-deficient Mre11 mutant (*mre11-3*) can be suppressed by the absence of the Ku complex and in an Exo1-dependent manner (Foster et al., 2011). This suggests that Mre11 counteracts the action of Ku at CPT-induced DSBs ends, which limits Exo1-dependent DNA resection. Similarly, although the combination of *mre11-3* with *rad3-102* is not lethal, it sensitizes the cells to UV exposure, a context in which the load of replication stress is enhanced in the *rad3-102* background (Moriel-Carretero & Aguilera, 2010a) (**Figure 1B**). Strikingly, the increased UV sensitivity of *rad3-102 mre11-3* compared to *rad3-102* can also be completely suppressed by the absence of Ku in an Exo1-dependent manner (**Figure 1B, EV1C**). These data show that Mre11, Ku and Exo1 play similar roles in DNA damage repair induced by *rad3-102* or CPT. Finally, *rad3-102* has been found lethal in combination with both *rad51*Δ and *pol32*Δ, leading to the proposal that DNA repair in the absence of Rad51 in these cells is backed up by the presence of the Pol32 non-essential subunit of the DNA polymerase ∂ (Moriel-Carretero & Aguilera, 2010a, Moriel-Carretero & Aguilera, 2010b). Remarkably, we found that the *rad51Δ pol32*Δ double mutant was more sensitive to CPT than the *rad51*Δ single mutant (**Figure 1C**), suggesting that, as in *rad3-102* cells, Pol32 also partially compensates the absence of Rad51 to cope with CPT-induced DNA damage.

**Figure 1.**
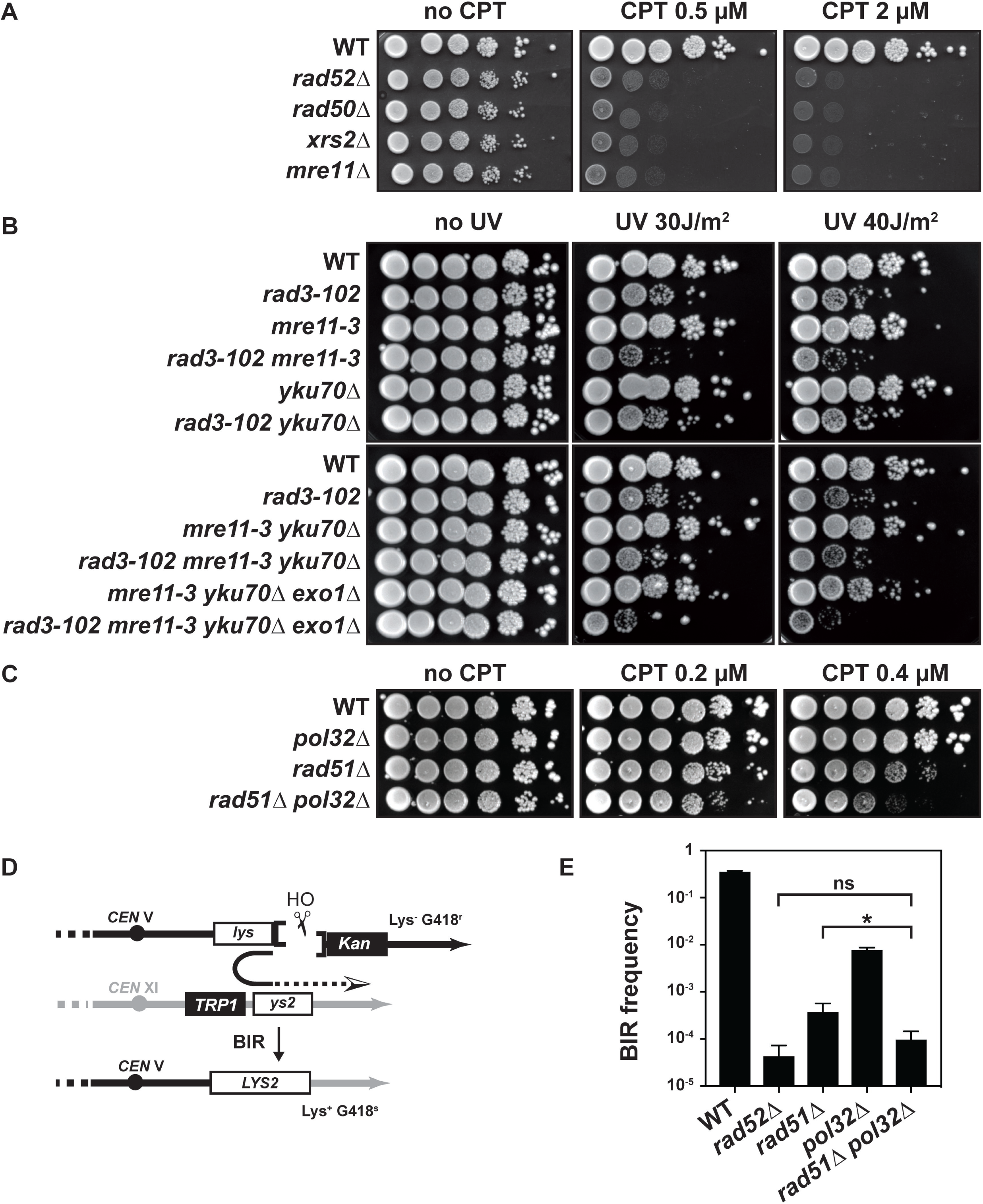
HR-mediated replication restart occurs by a BIR-like mechanism. **(A) (C)** CPT sensitivity assayed by 10-fold serial dilutions of different mutant combinations on YPAD plates. **(B)** UV sensitivity assayed by 10-fold serial dilutions of different mutant combinations with *rad3-102* allele on YPAD plates after exposure to the indicated UV-C doses. **(D)** Schematic representation of the BIR assay. Two fragments of the *LYS2* gene sharing 2.1 kb of homology (*lys* and *ys2*) were integrated on chromosomes V and XI, respectively. Induction of *HO* endonuclease expression under control of the *GAL1* promoter produces a DSB next to the *lys* fragment that can be repaired by BIR. BIR events can be scored by selecting survivor colonies, which harbor a functional *LYS2* gene. **(E)** BIR frequencies (Lys+ survivors among total cells) for the WT and indicated mutant strains are plotted on a logarithmic scale. Data represent the mean ± SD from 3 to 6 independent experiments. ns, non-significant. * *P*-value = 0.0134; Mann-Whitney unpaired t test.

Thus, *rad3-102* cells suffer from a replication stress that mimics the effect of CPT. However, accumulation of topological stress is not expected in *rad3-102* cells, since they are not affected in DNA supercoils removal by Top1. Tdp1 (Tyrosyl-DNA phosphodiesterase 1) is not required for survival in *rad3-102* cells, consistent with the TFIIH complex not being covalently bound to DNA through a tyrosine residue. Interestingly, however, the absence of the metalloprotease Wss1, which clears chromatin-bound sumoylated proteins in response to genotoxic stress (Balakirev et al., 2015), impacts the sensitivity of *rad3-102* cells to UV (**Figure EV1B**). This suggests that, as for the removal of Top1 in CPT-treated cells, Wss1 is involved in the removal of the compromised TFIIH complex in *rad3-102* cells. Overall, in view of the strikingly similar genetic requirements for the cell survival in response to CPT and in the *rad3-102* background, we decided to use these two systems to further investigate the repair mechanisms required to cope with this type of replication stress, independently from the accumulation of DNA supercoiling and DNA-protein crosslinks.

### Rad51 and Pol32 independently promote DSB repair by break-induced replication

First, we asked what could be the specific contribution of Pol32 to DNA repair in the absence of Rad51. We previously proposed that Pol32 may stabilize strand invasion in the absence of Rad51 by promoting the priming of DNA synthesis (Moriel-Carretero & Aguilera, 2010b). Among DSB repair pathways mediated by HR, Pol32 has only an essential role during BIR. In the absence of Pol32, the initiation of DNA synthesis during BIR is compromised (Lydeard et al., 2007). Moreover, the recovery of both Rad51-dependent and Rad51-independent survivors I cells lacking telomerase, thought to occur by BIR, also rely on Pol32 (Lydeard et al., 2007). Finally, the BIR pathway has been proposed to be involved in the repair of broken replication forks (Mayle et al., 2015). We thus wondered if Pol32 could help Rad51 and compensate for its absence to promote BIR in cells exposed to CPT or in the *rad3-102* background. To assess the redundant role of Pol32 over Rad51 in BIR, we used a well-described chromosomal system (Donnianni & Symington, 2013) in which a single DSB is induced by the HO endonuclease. In this system, only one of the two ends can undergo homology-dependent strand invasion at an ectopic location. Subsequent priming and elongation of DNA synthesis reaching the chromosome end leads to the production of viable Lys2+ recombinants (**Figure 1D**). In this system, the absence of Rad52 decreased the BIR frequency by about three orders of magnitude compared to the wild type (**Figure 1E**). Deletion of *RAD51* or *POL32* also significantly decreased the BIR frequency compared to the wild type (**Figure 1E**) (Donnianni & Symington, 2013). However, only the combined absence of Rad51 and Pol32 did affect the BIR frequency as much as in the absence of Rad52 (**Figure 1E**). These results show that BIR mainly occurs by a Rad52- and Rad51-dependent pathway, but that Pol32 can also promote BIR in the absence of Rad51. Because we observed that Pol32 promotes cell survival in the absence of Rad51 both in response to CPT and in *rad3-102* cells, we propose that DNA repair in these contexts occurs by a BIR-like mechanism.

### Mus81 is involved in HR-mediated repair involving both Rad51 and Pol32

In human cells, the structure-specific endonuclease (SSE) Mus81 has been shown to generate DSBs in response to CPT treatment, leading to the suggestion that Mus81 may cleave stalled or reversed replication forks upon Top1 poisoning to promote fork restart (Regairaz et al., 2011). In budding yeast, null and catalytically dead (*mus81-dd*) mutations of *MUS81* have been shown to sensitize cells to mild CPT doses (**Figure 2A, EV1A**) (Bastin-Shanower et al., 2003, Blanco et al., 2010, Liu et al., 2002, Tay & Wu, 2010, Vance & Wilson, 2002). The absence of Mus81 is backed up by the Yen1 SSE, as shown by the higher CPT sensitivity of the double *mus81*Δ *yen1*Δ mutant compared to *mus81*Δ (**Figure EV2A**) (Blanco et al., 2010, Tay & Wu, 2010). Moreover, Mus81 and Yen1 have been proposed to limit BIR synthesis initiated from a replication-born DSB (Mayle et al., 2015). We asked if Mus81 could have a similar role in CPT-induced DNA damage repair. First, we observed that the combination of *mus81*Δ with *rad51*Δ and *pol32*Δ mutations resulted in an increased CPT sensitivity than either single mutants (**Figure 2B**) (Mayle et al., 2015), suggesting that Mus81 participates to both Rad51- and Pol32-dependent repair.

**Figure 2.**
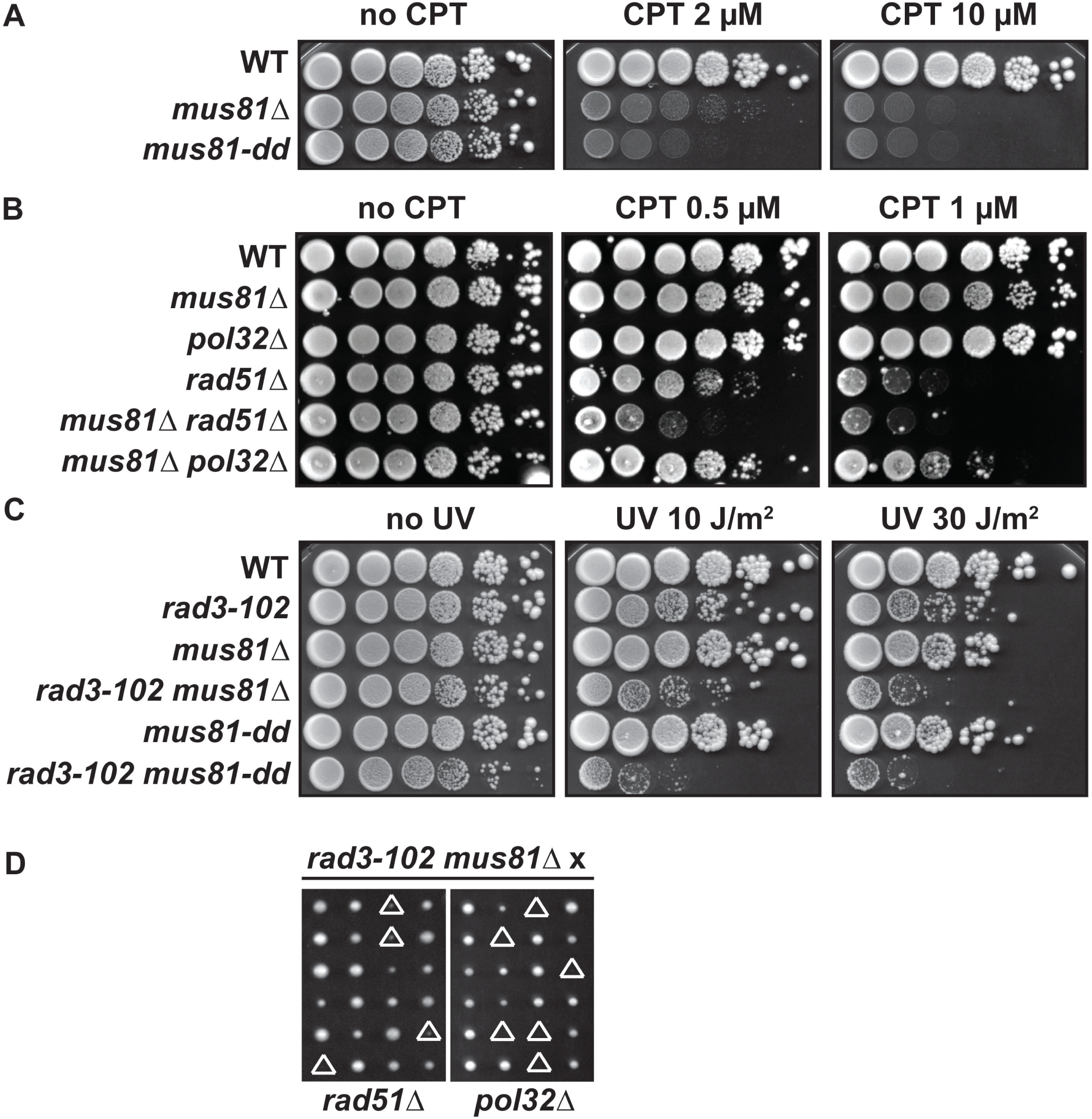
Mus81 is required in both Rad51- and Pol32-dependent restart pathways. **(A) (B)** CPT sensitivity assayed by 10-fold serial dilutions of different mutant combinations on YPAD plates. **(C)** UV sensitivity assayed by 10-fold serial dilutions of different mutant combinations with *rad3-102* allele on YPAD plates after exposure to the indicated UV-C doses. **(D)** Synthetic combinations of *rad3-102 mus81*Δ with *rad51*Δ and *pol32*Δ. Tetrads dissected on YPAD medium are shown. Triangles indicate either a severe growth defect or lethality.

As for *rad51*Δ, *mus81*Δ or *mus81-dd* are not lethal in combination with *rad3-102* but the double mutant cells exhibited a much higher sensitivity to UV than single mutant cells (**Figure 2C**) (Moriel-Carretero & Aguilera, 2010a). We analyzed the redundancy between Mus81 and Yen1 for survival in the *rad3-102* background. *rad3-102 yen1*Δ cells were not more sensitive to UV than *rad3-102* cells (**Figure EV2B**), consistent with *yen1*Δ cells not being sensitive to CPT (**Figure EV2B)**. The *rad3-102 mus81*Δ *yen1*Δ triple mutants were unviable (**Figure EV2C**), indicating that Yen1 also backs up Mus81 in *rad3-102* cells. Finally, we found that *rad3-102 mus81*Δ *pol32*Δ mutants were unviable and *rad3-102 mus81*Δ *rad51*Δ cells had a severe growth defect (**Figure 2D**). These results indicate that Mus81 is required for repair mediated by both Rad51 and the polymerase δ subunit Pol32 in *rad3-102* cells.

### Mus81 is not required for replication progression upon Top1 poisoning

Next, we reasoned that if an increased CPT sensitivity is associated with replication defects, we should observe an impaired progression through S phase. Opposite to other cellular models, it was reported that an acute exposure to CPT in liquid cultures of *S. cerevisiae* cells did not induce a delay in S phase progression but rather a prolonged arrest in G_2_/M (Puddu et al., 2017, Ray Chaudhuri et al., 2012, Redon et al., 2003). CPT is highly insoluble in culture media and the water-soluble derivatives of CPT (topotecan and irinotecan) do not affect yeast cells growth nor their survival (Del Poeta et al., 1999), thus not being exploitable for our study. We suspected a permeability issue for CPT entry into the cells, as described for the proteasome inhibitor MG132 and the DNA polymerase inhibitor Aphidicolin (Liu et al., 2007, Plevani et al., 1980). We therefore used a modified culture medium described to render cells more permeable to the antifungal agent brefeldin A (Pannunzio et al., 2004) in order to increase the cell permeability to CPT. We took advantage of the natural blue fluorescence emitted by the CPT compound when excited by 350 nm UV light to quantify the relative amount of CPT inside cells. Incubation of an asynchronous cell culture with CPT for 30 minutes in MPD +SDS medium (minimal-proline-dextrose + 0.003% SDS) (Pannunzio et al., 2004) allowed the detection of blue fluorescence inside cells, whose quantification was up to 45 times higher than in control cells incubated with DMSO in the same medium or in cells incubated with CPT for 30 minutes in standard MAD medium (minimal-ammonium-dextrose) **(Figure EV3A,B**). We then used these conditions to analyze the progression through a single S phase in the presence of CPT by flow cytometry. Wild type cells were synchronized in G_1_ with α-factor and incubated with CPT for 1 hour, then released from G_1_ into S phase, still in the presence of CPT. As observed by others, CPT did not alter S phase progression when cells were grown in normal MAD medium (**Figure EV3C**) (Puddu et al., 2017, Ray Chaudhuri et al., 2012, Redon et al., 2003), but when incubated with CPT in MPD +SDS medium, they progressed significantly slower through S phase than control wild type cells incubated with DMSO, thus reaching the G_2_ phase later (**Figure EV3C**). These experimental conditions allowed us for the first time to characterize the effect of CPT on DNA replication using *S. cerevisiae* cells as a model system.

Using these conditions, we next asked how the HR factors Rad52 and Rad51 and the Mus81 resolvase contribute to the progression of replication forks encountering poised Top1. At the global level, using flow cytometry, we could observe that the S phase delay was strikingly increased in cells lacking one of the two main HR factors, Rad52 or Rad51, compared to wild type cells (**Figure 3A**). However, in the absence of Mus81, S phase progression was as affected by CPT as in wild type cells. We confirmed these results using the *rad3-102* mutation. We synchronized cells in G_1_ phase with α-factor and irradiated them with UV-C. Cells were kept for 2 hours in G_1_ before release into S phase, in order to let compromised NER leave DNA gaps or nicks bound by the TFIIH complex in the *rad3-102* mutant, as described previously (Moriel-Carretero & Aguilera, 2010a). As expected, UV irradiation only affected S phase progression in *rad3-102* cells and to a greater extent in *rad3-102* cells lacking the HR factor Rad51 (**Figure 3B**). However, *rad3-102* cells lacking Mus81 or Mus81 nuclease activity did not show an increased S phase delay compared with *rad3-102* single mutants (**Figure 3C**). These results indicate that HR factors Rad52 and Rad51 but not Mus81 are required to promote S phase progression when replication stress is induced by CPT.

**Figure 3.**
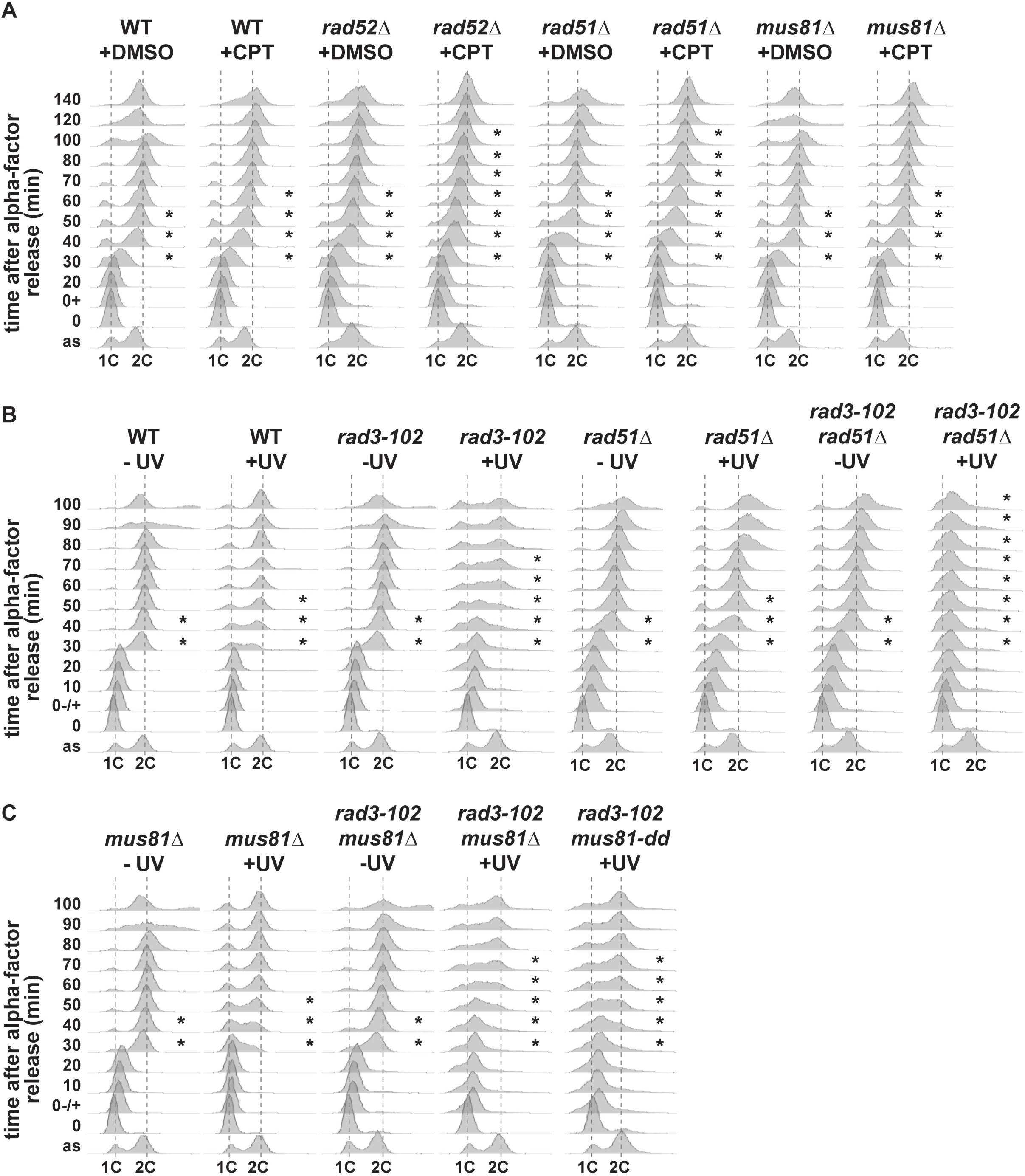
Mus81 is not required for S phase progression in CPT-treated and *rad3-102* cells. **(A)** Analysis of DNA content by flow cytometry of G_1_ phase synchronized wild type, *rad52*Δ, *rad51*Δ, *mus81*Δ and *mus81-dd* cells and further released into S phase. Cells were synchronized in G_1_ with α-factor, treated with DMSO or 100 µM CPT, let in G_1_ for 1 h, and released into S phase. **(B)** Analysis of DNA content by flow cytometry of G_1_ phase synchronized wild type, *rad3-102*, *rad3-102 rad51*Δ, *rad3-102 mus81* and *rad3-102 mms4*Δ cells and further released into S phase. Cells were synchronized in G_1_ with α-factor, untreated or irradiated with 20 J/m^2^ UV-C, let in G_1_ for 2h, and released into S phase. Asterisks indicate the progression of cells in S phase.

To confirm these data at the molecular level, we analyzed S phase progression by DNA combing from asynchronous cell cultures treated with CPT. DNA combing allows monitoring replication fork progression genome-wide though at the level of individual DNA molecules (Bianco et al., 2012, Tourrière et al., 2017). After incubation with CPT for 2 hours, cells were pulse-labeled with the thymidine analog EdU for 20 minutes before combing analysis (**Figure 4A**). We first compared the length of the EdU tracks in all strains under unchallenged conditions (+DMSO). Only *mus81*Δ cells showed a significant decrease of the EdU track length compared to wild type cells (**Figure 4A**; *P*-value = 0.0015). This result agrees with a recent report that proposed a role for human Mus81 in DNA replication in the absence of exogenous damage (Fu et al., 2015). Then, we focused our attention onto the effect of CPT on fork progression in each strain. As expected from cell cycle analyses (**Figure 3A**), CPT treatment significantly affected fork progression in all strains compared to the DMSO control (*P*-value < 0.0001) (**Figure 4A**). The EdU track length of wild type cells was reduced by 47% and that of mutant cells was further reduced to 55% in *rad51*Δ cells and 75% in *rad52*Δ cells (**Figure 4A**). However, the reduction in fork progression caused by CPT in *mus81*Δ cells (45%) was similar to the one of wild type cells (**Figure 4A**). This was confirmed in *rad3-102 mus81*Δ cells, in which replication fork progression assayed by DNA combing was not more affected than in single mutants (**Figure EV4**). Overall, our results show that CPT-mediated Top1 poisoning induces a global replication stress that requires HR factors for replication fork restart, but argue against a role of Mus81 in restart, as was suggested in human cells (Regairaz et al., 2011).

**Figure 4.**
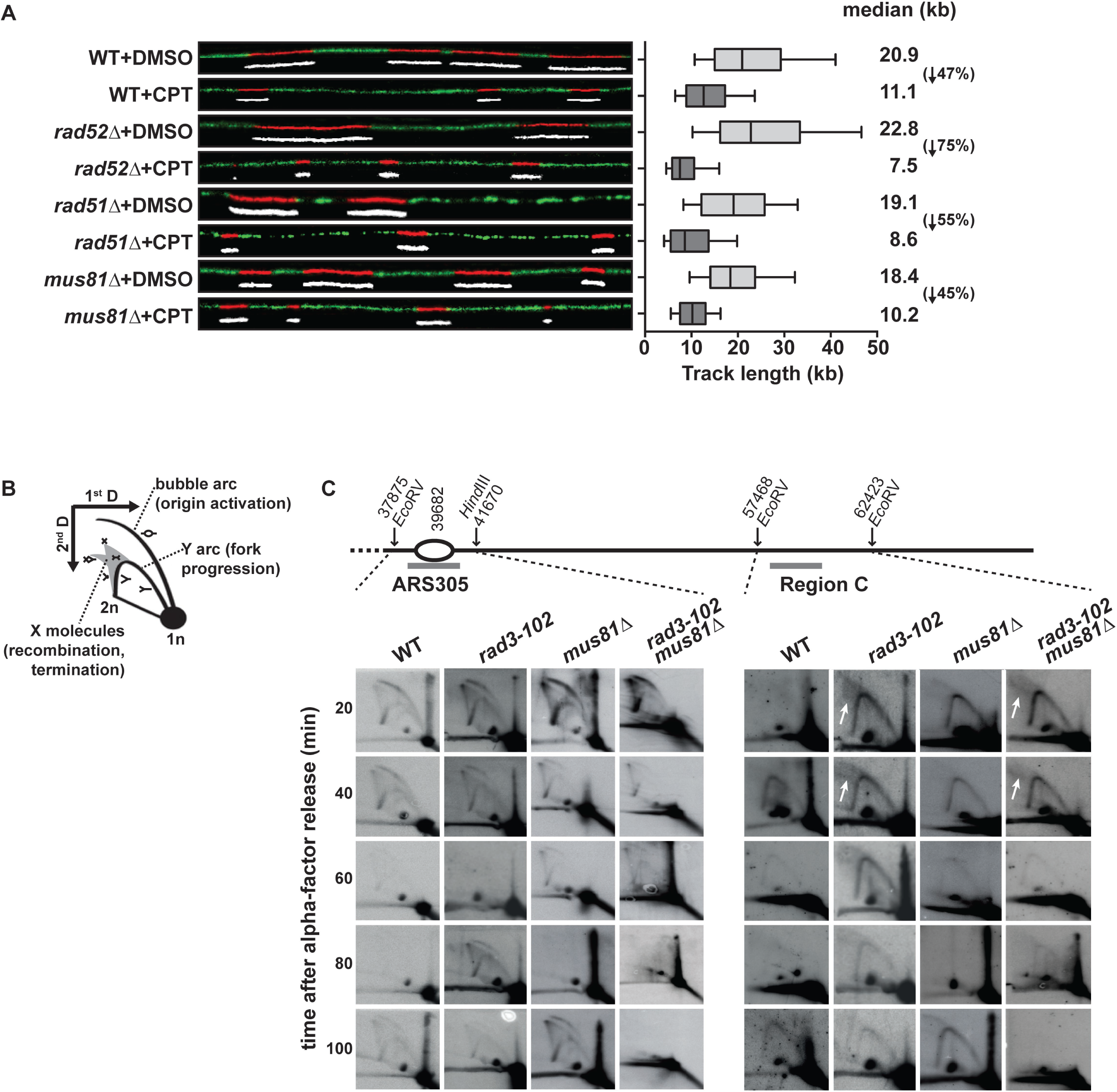
Rad52 and Rad51 are required for HR-mediated replication progression but not Mus81. **(A)** Analysis of replicated DNA tracks length by single-molecule DNA combing in WT, *rad52*Δ, *rad51*Δ and *mus81*Δ cells exposed to CPT. Exponentially growing cells were treated with DMSO or 50 µM CPT for 2h and then pulse-labeled with 50 µM EdU for 20min. DNA fibers were combed on silanized coverslips and EdU-labeled DNA was detected by Click chemistry. Graph depicts the distribution of EdU tracks length in kb. Box and whiskers indicate 25-75 and 10-90 percentiles, respectively. Median EdU tracks length is indicated in kb. The decrease percentage in EdU tracks lengths between the DMSO and CPT conditions is indicated between parentheses for each strain. Representative images of DNA fibers are shown. Red and white: EdU, green: DNA. **(B)** Schematic representation of replication intermediates visualized by 2D gels. **(C)** Analysis of replication intermediates by 2D gel electrophoresis. Replication intermediates were monitored at early origin *ARS305* and region C in WT, *rad3-102*, *mus81*Δ and *rad3-102 mus81*Δ cells. Cells were synchronized in G_1_ with α-factor and collected at the indicated time points after release into S phase. A scheme of the studied chromosomal region is shown (drawn to scale). Relevant probes are indicated by gray bars, and coordinates of ARS and restriction sites are indicated in bp. Accumulation of recombination molecules is indicated by white arrows.

### Mus81 resolves S phase-induced recombination events in G_2_/M

Our data indicate that Mus81 is required to cope with the replication stress induced by Top1 poisoning but it does not promote replication progression in S phase. In human cells, Mus81 has been proposed to promote fork restart by cleaving stalled or reversed replication forks (Regairaz et al., 2011). To understand this apparent contradiction and define the precise window of Mus81 activity, we assessed replication progression by two-dimensional (2D) neutral-neutral gel electrophoresis in synchronized cultures after release from G_1_ phase, which was previously used to characterize the replication defects in *rad3-102* cells. We studied the early replication origin *ARS305* and the passively replicated region C besides it (Lopes et al., 2003). Notably, *rad3-102* cells accumulated complex branched structures at region C (**Figure 4B,C**, see arrows), described as recombination or fork reversal events (Moriel-Carretero & Aguilera, 2010a). This assay gave us the opportunity to study the role of Mus81 in replication fork progression and recombination intermediate resolution in the *rad3-102* background. As previously described, firing at *ARS305* occurs slightly earlier in *rad3-102* cells compared to wild type (Moriel-Carretero & Aguilera, 2010a). In the absence of Mus81 (*mus81*Δ and *rad3-102 mus81*Δ mutants), slower replication was observed around the *ARS305*, as the Y arc signal was still clearly observable at 60 minutes, while it had already disappeared in wild type and *rad3-102* cells (**Figure 4C**, left panel). This is consistent with a slower replication fork progression in *mus81*Δ than in wild type cells observed in the absence of CPT-induced DNA damage (**Figure 4A, EV4**). We could not observe an accumulation of recombination intermediates *rad3-102 mus81*Δ cells compared to *rad3-102* cells (**Figure 4C**, see arrows), suggesting that Mus81 does not process complex branched structures observed in *rad3-102* cells. More interestingly, we noted that, in a 100 minutes time-window after G_1_ release, *ARS305* fired twice in wild type, *mus81*Δ and *rad3-102* cells, implying two rounds of replication. This was not observed in the *rad3-102 mus81*Δ mutant (**Figure 4C**, left panel). As for *ARS305* region, replication forks progressed only once through region C in *rad3-102 mus81*Δ cells (**Figure 4C**, right panel). These results made us consider a cell cycle delay that could stem from a replication termination defect in the absence of Mus81 in *rad3-102* cells.

To explore this, we monitored the appearance of YFP-Rad52 foci in cells exposed to CPT. Wild type cells were synchronized in G_1_ with α-factor and incubated with CPT for 30 minutes, then released from G_1_ into S phase in the presence of CPT. Wild type cells incubated with DMSO showed the appearance of a low amount of Rad52 foci in S phase 40 and 60 minutes after release from G_1_ (**Figure 5A**). These foci disappeared when cells reached G_2_/M (**Figure 5A**), consistent with the observation that spontaneous Rad52 foci predominantly form in S phase cells (Lisby et al., 2001). When wild type cells were released from the G_1_ arrest in the presence of CPT, they accumulated 6 times more Rad52 foci than in control cells only when cells entered S phase at 40 minutes (**Figure 5A**). This accumulation continuously increased until cells reached the G_2_/M phase at 80 minutes and then started to decrease in later time points (**Figure 5A**). These results show that S phase entry is required for the initiation of recombination events induced by CPT and these events are resolved after the completion of DNA replication in G_2_/M. When assessing the contribution of Mus81, we made two main observations. First, the increased accumulation of Rad52 foci during S phase caused by CPT exposure was not suppressed in the absence of Mus81 (**Figure 5B**). This confirmed that Mus81 is not required during S phase to process recombination intermediates (**Figure 4B**). Second, we observed that CPT-induced Rad52 foci in *mus81*Δ cells accumulated over the entire time course experiment until 120 minutes, not showing the decrease observed in wild type cells during the G_2_/M phase (**Figure 5A,B**). This phenotype was again confirmed in *rad3-102 mus81*Δ cells, which accumulated more Rad52 foci than either single mutant (**Figure EV5A**). These results suggest that Mus81 processes S phase-induced recombination events in G_2_/M.

**Figure 5.**
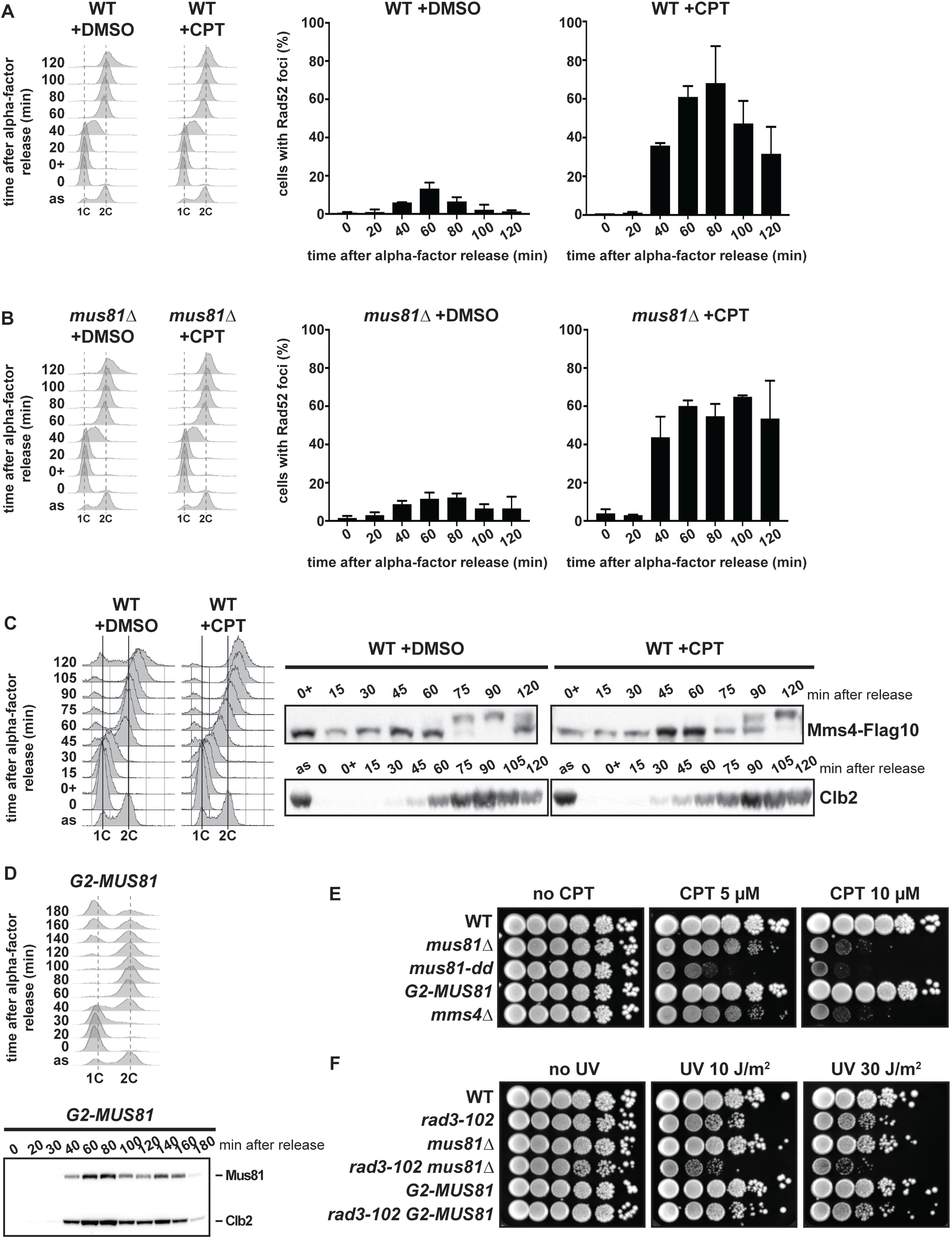
Mus81 is involved in the processing of recombination intermediates in G_2_/M after replication restart. **(A) (B)** Kinetic analysis of Rad52 foci formation. Wild-type and *mus81*Δ cells containing Rad52-YFP and mCHERRY-Pus1 were synchronized in G_1_ with α-factor, treated with DMSO or 50 µM CPT, let in G_1_ for 30 min, and released into S phase. Cells were collected at the indicated time points and visualized by fluorescence microscopy. Mean and SD of cells with Rad52 foci for three independent experiments are shown for each time point. An illustrative image is shown with the nuclear compartment marker Pus1 in red and Rad52 foci in yellow. Flow cytometry profiles corresponding the experimental setup are shown. **(C)** Mms4-Flag10 phosphorylation analyzed by immunoblot in wild type cells exposed to CPT. Wild-type cells were synchronized in G_1_ with α-factor, treated with DMSO or 50 µM CPT, let in G_1_ for 1 h, and released into S phase. Cells were collected at the indicated time points and Mms4 was immunodetected with Flag antibodies. Clb2 immunodetection serves as a marker for G_2_ phase entry. FACS profiles corresponding the experimental setup are also shown. **(D)** Immunoblot analysis of G2-Mus81. *G2-MUS81* cells were synchronized in G_1_ with α-factor and released into S phase. Cells were collected at the indicated time points and G2-Mus81 and Clb2 were immunodetected with Clb2 antibodies. Flow cytometry profiles corresponding the experimental setup are shown. **(E)** CPT sensitivity assayed by 10-fold serial dilutions of *G2-MUS81* compared to the wild type, *mus81* and *mms4*Δ mutants. **(F)** UV sensitivity assayed by 10-fold serial dilutions of *G2-MUS81* in combination with *rad3-102* allele compared to *rad3-102 mus81*Δ on YPAD plates after exposure to the indicated UV-C doses.

In an unperturbed cell cycle, the regulating subunit Mms4 of Mus81-Mms4 complex is hyper-phosphorylated by multiple kinases at the G_2_/M transition, and this correlates with an enhanced activity of the complex on branched DNA structures *in vitro* (Gallo-Fernandez et al., 2012, Matos et al., 2011, Matos et al., 2013, Princz et al., 2017). To validate if the temporal regulation of Mus81 activity fits with the timely requirement for Mus81 to process recombination intermediates generated during CPT-induced damage repair (**Figure 5B**), we followed a synchronous culture going through a single S phase and analyzed Mms4 phosphorylation. Treatment of wild type cells with CPT caused a delay in S phase progression with respect to control cells, with a consequent delay in Mms4 phosphorylation, detected as an electrophoretic mobility shift by immunoblotting (**Figure 5C**). The highest degree of Mms4 phosphorylation was reached 90 and 120 minutes after G_1_ release in control cells and cells treated with CPT, respectively (**Figure 5C**). In *rad52*Δ cells exposed to CPT, cells arrested in late S phase and no phosphorylation of Mms4 was observed (**Figure EV5B**). We also analyzed Mms4 phosphorylation in wild type and *rad3-102* cells. First, it is worth noting that exposure to of UV-C in G_1_ phase did not induce any change of Mms4 mobility in both tested strains (**Figure EV5C**). In wild type cells, Mms4 underwent phosphorylation 60 to 70 minutes after release from G_1_, when most cells had reached the G_2_ phase. However, no phosphorylation of Mms4 was observed in *rad3-102* cells, in agreement with their accumulation in S phase after G_1_ release (**Figure EV5C**). Thus, the nuclease activity of Mus81-Mms4 is required to process S phase-associated HR events once the cells reach G_2_ (validated by the accumulation of cyclin B2 (Clb2)) (**Figure 5C**). Indeed, hyper-phosphorylation of Mms4 is required for the function of Mus81 in the repair of CPT-induced DNA damage, as the *mms4-14A* mutant, which cannot undergo phosphorylation by Cdk1 and Cdc5 (Matos et al., 2011) is sensitive to CPT (**Figure EV5D**). However, the CPT sensitivity of the *mms4-14A* mutant was lower than in the complete absence of Mms4 (*mms4*Δ), and the *mms4-9A* mutant (*mms4-np*, (Gallo-Fernandez et al., 2012)) was not found more CPT sensitive than the wild type (**Figure EV5D**). This discrepancy between *mms4* mutants could be explained by the high number and redundancy of Mms4 phosphorylation sites (Matos et al., 2011, Princz et al., 2017). Last, to confirm the need of Mus81 only after the completion of genome duplication, we limited the expression of Mus81 to the G_2_/M phase by taking advantage of the regulatory elements of the mitotic cyclin B2 (Karras & Jentsch, 2010). Fused to this G_2_ tag, Mus81 was only expressed in cells once cells reached the G_2_ phase and was targeted for degradation in the following G_1_ phase (**Figure 5D**). Contrary to the cells lacking Mus81, Mms4 or the catalytic activity of Mus81 (*mus81-dd*), cells bearing the *G2-MUS81* allele were not sensitive to CPT-induced DNA damage (**Figure 5E**). Analogously, *rad3-102 G2-MUS81* cells were not more sensitive to UV than the *rad3-102* single mutant (**Figure 5F**).

Overall, we conclude that HR factors are required for the restart of replication forks blocked by Top1 poisoning by mediating recombination events. Despite that these events occur in S phase, Mus81 nuclease is acting to process recombination events only when its activity is increased during the G_2_/M phase. Our results also show that Mus81 was not required for the assembly of Rad52 foci, nor was activated at the time of their appearance, indicating that Mus81 is unlikely to be involved in the generation of HR substrates by cleaving replication forks in our systems.

### Mus81 processes recombination intermediates to complete replication replication

Our 2D-gel analysis indicated that *rad3-102 mus81*Δ cells synchronously released from a G_1_ block had a delay in starting the following cell cycle (**Figure 4C**). We confirmed this observation in cells exposed to Top1 poisoning by performing longer time course flow cytometry experiments. Indeed, *mus81*Δ cells treated with CPT started the following cell cycle 40 minutes later than wild type cells under the same treatment (**Figure EV6A**). This cell cycle delay could stem from the inability of cells to timely process recombination intermediates required for the restart of replication. Since we proposed that the repair of CPT-induced DNA damage could occur through a BIR-like mechanism, this implies that replication restart should occur by a migrating D-loop. Merging of the D-loop with a converging replication fork would form a single Holliday junction, whose resolution would be mandatory for replication termination. We did not observe an accumulation of recombination or termination intermediates in *rad3-102 mus81*Δ cells, yet they may have not accumulated in the region analysed by 2D-gels. Thus, we decided to look for the accumulation of termination intermediates at the genome level by pulsed-field gel electrophoresis (PFGE). PFGE allows the separation of individual chromosomes according to their size in G_1_ and G_2_/M cells. However, when cells enter S phase, the chromosomes are trapped into the wells due to the presence of joint molecules (JMs), as replication bubbles and other replication intermediates. For instance, this could be observed for wild type or *mus81*Δ cells incubated with DMSO, for which chromosome bands disappeared from the gel 40 minutes after the release from G_1_ and could be detected back again at 60 minutes (**Figure 6B**, upper panel). After Southern blot analysis, we could observe that chromosome IV was retained in the gel well at S-phase entry 40 minutes after release and then progressively disappeared from the wells (**Figure 6B**, lower panel). This is consistent with the FACS analysis, which shows that bulk DNA synthesis started at 40 minutes and ended 60 to 80 minutes after release (**Figure 6A**). Incubation of wild type cells with CPT induced a delay in S phase progression (**Figure 6A**), which was mirrored by the kinetics of accumulation of JMs in the gel wells (**Figure 6B**). As shown before (**Figure 3A**), CPT treatment induced the same S phase progression delay in *mus81*Δ and wild type cells (**Figure 6A**). Yet, opposite to the situation in wild type cells, the amount of JMs did not decrease 100 minutes after release from G_1_ in CPT-treated *mus81*Δ cells and chromosomes hardly re-entered the gel (**Figure 6B**). Since the bulk of DNA synthesis had already ended at that time, these JMs unlikely represents replication forks but rather single Holliday junctions that would form upon the merging of D-loops with converging forks. Therefore, these results suggest that Mus81 would be required for the processing of termination intermediates specifically formed during the restart of replication caused upon Top1 poisoning.

**Figure 6.**
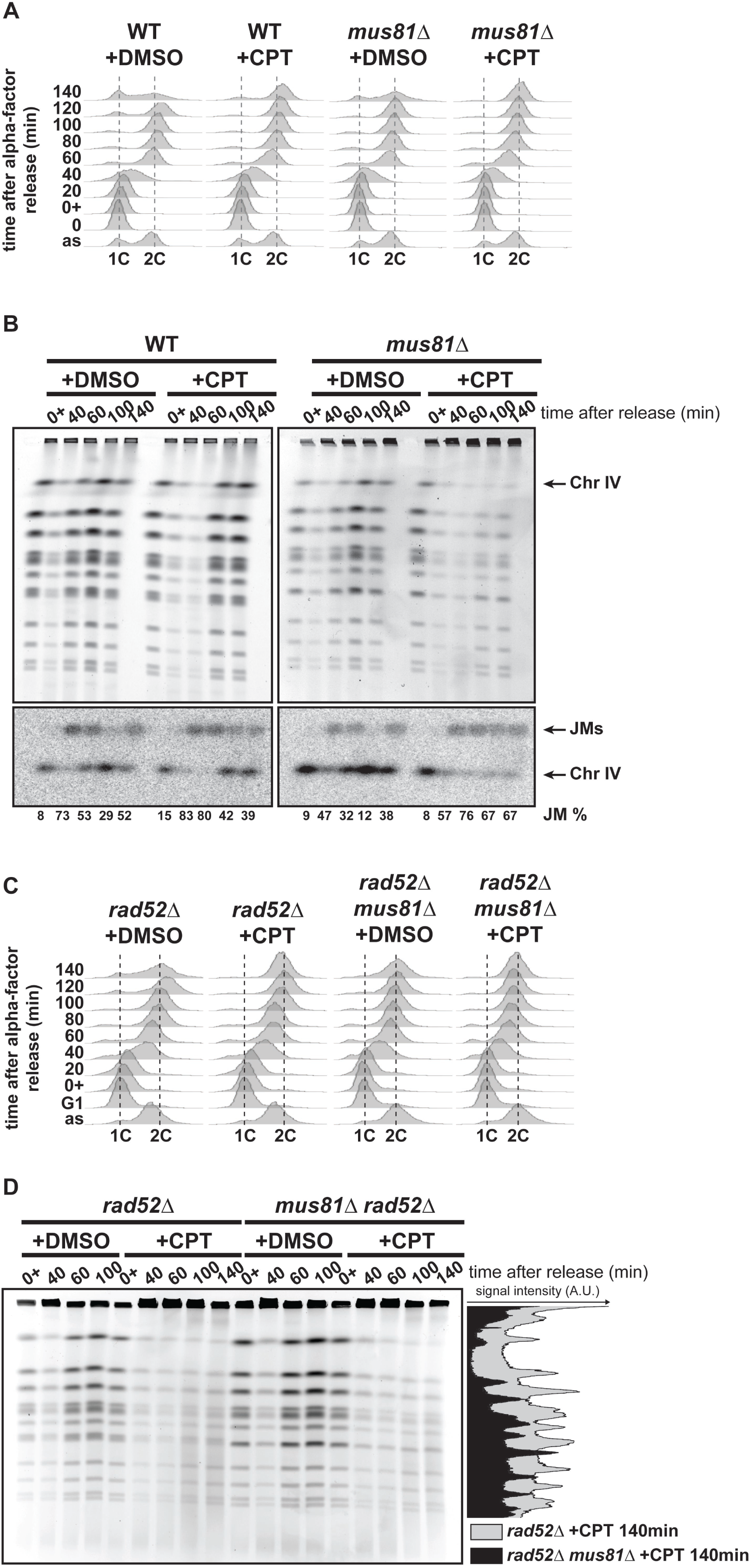
Recombination intermediates generated by replication restart accumulate in the absence of Mus81. **(A) (B)** Pulsed-field gel electrophoresis (PFGE) analysis of wild type and *mus81*Δ cells in response to CPT. Wild type and *mus81*Δ cells were synchronized in G_1_ with α-factor, treated with DMSO or 100 µM CPT, let in G_1_ for 1 h, and released into S phase. Cells were collected at the indicated time points. DNA contents was analyzed by flow cytometry and the DNA extracted in agarose plugs was analyzed by PFGE. Upper panel: agarose gel stained with ethidium bromide; lower panel: Southern blot using a chromosome IV specific probe. JMs, joint molecules accumulated in the gel wells. Quantification of JMs relative to the total amount of DNA is indicated for each time point. **(C) (D)** PFGE analysis of *rad52*Δ and *rad52*Δ *mus81*Δ cells in response to CPT performed as in (A) (B). The gel has been stained with Ethidium bromide and densitometry profiles corresponding to the +CPT 140 min time points in *rad52*Δ and *rad52*Δ *mus81*Δ cells are shown.

One last remaining question was whether this BIR-mediated replication restart occurs from broken or reversed replication forks. The stabilization of a nicked DNA bound to Top1 by CPT has been initially proposed to induce a replication fork run-off, leading to the formation of a one-ended DSB (Strumberg et al., 2000). More recently, an alternative model has been proposed, integrating that Top1 inhibition by CPT causes the accumulation of topological constrains in DNA that may impede the progression of replication forks (Koster et al., 2007). Since 20 to 25% of forks have been observed to be reversed by electron microscopy after *in vivo* psoralen crosslinking in CPT-treated yeast cells, it was suggested that the accumulation of supercoils in front of replication forks promote their reversal (Menin et al., 2018, Ray Chaudhuri et al., 2012). Reversed forks are formed upon the annealing of nascent strands together, exposing a single double-strand DNA end that mimics a one-ended DSB. In PFGE experiments, the occurrence of DSBs would result in the diminution of full-length chromosome bands and the appearance of smeared signals. This could be readily observed wild type cells released in S phase in the presence of the radiomimetic agent zeocin, which induces fragmentation of chromosomes (**Figure EV6B**). We could not observe such smears in wild type CPT-treated cells when they progressed through S phase 30 to 60 minutes after release from the G_1_ block (**Figure 6B, EV6B**). Similarly, we could not observe any smeared signal in S-phase *rad3-102* cells (**Figure EV6C**). In the absence of Rad52, which is strictly required for HR-mediated repair of DSBs, smears could be detected only when CPT-treated cells reached the G_2_ phase 100 minutes after release (**Figure 6C,D**). This could be explained by the accumulation of unrepaired CPT-induced DSBs that could not be detected in S phase. Alternatively, the appearance of broken chromosomes in G_2_ *rad52*Δ cells could be the consequence of the cleavage of blocked or reversed forks that could not undergo restart through HR. Indeed, densitometry analysis of PFGE showed that the amount of smeared signals at 140 minutes post-release in *rad52*Δ cells treated with CPT was reduced by 50% in the absence of Mus81 (**Figure 6C,D**).

Together, our results favour a model in which Top1 poisoning by CPT does not induce DSBs during S phase. Mus81 is required to process recombination intermediates that accumulate at termination sites following HR-mediated replication forks restart and forks that could not be restarted because of a HR defect.

## DISCUSSION

In the present study, we have used Top1 poisoning by CPT to generate a replication stress genome-wide and to study the restart replication forks blocked by Top1ccs. Our results support a model for replication restart mediated by HR through a BIR-like mechanism occurring in S phase. Replication progression primarily depends on Rad52 and Rad51. The Mus81 nuclease does not participate to the replication restart but appears to be essential to resolve recombination intermediates to promote the termination of those restarted forks. Thanks to a complementary, though independent, approach using the *rad3-102* allele, we conclude that this mechanism is independent of the accumulation of DNA supercoiling and DNA-protein crosslinks naturally caused by CPT.

It is worth emphasizing that this work has been made possible because we set up conditions to improve CPT entry into yeast cells in liquid cultures. Using standard culture conditions, CPT treatment did not detectably delay S phase completion, even in *rad52*Δ mutants, whereas this phenotype was clearly observed in CPT-treated human cells (Ray Chaudhuri et al., 2012). The culture conditions we used allowed observing a CPT-dependent S phase delay, which could be exacerbated by mutants defective in HR as *rad52*Δ and *rad51*Δ (**Figure 3**). These S phase progression defects could be characterized at the molecular level by demonstrating replication defects by both DNA combing and pulsed-field gel electrophoresis (PFGE) experiments (**Figure 4** and **Figure 6**). The latter experiments could also show physical evidence of chromosome breakage upon CPT treatment, which were not found in previous studies (Ray Chaudhuri et al., 2012, Redon et al., 2003). Nevertheless, DSBs only appeared in CPT-treated *rad52*Δ cells and not during their progression through S phase, as expected from the conversions of nicked DNA into DSBs by the passage of replication forks (Hsiang et al., 1989, Nitiss & Wang, 1988, Strumberg et al., 2000). DSBs appeared when cells reached the G_2_ phase and were partially dependent on the presence of the Mus81 nuclease (**Figure 6**). The remaining DSBs are expected to be dependent on the Yen1 nuclease because the survival of *mus81*Δ cells is sustained by the presence of Yen1 in response to CPT and in *rad3-102* cells (**Figure EV2A,C**) (Blanco et al., 2010, Tay & Wu, 2010). Consistently, Yen1 catalytic activity is also restricted to G_2_/M (Blanco et al., 2014, Eissler et al., 2014, Matos et al., 2011).

CPT-induced DSBs were also detected in human cells and partially Mus81-dependent (Regairaz et al., 2011). In these cells, immunostaining of γH2AX, a marker of DNA damage checkpoint activation, was co-localizing with sites of neo-synthesized DNA. This led the authors to propose that Mus81 cleaves stalled replication forks to promote their restart (Regairaz et al., 2011). However, these experiments were performed in HCT116 colorectal carcinoma cells, which are deficient for Mre11 (Giannini et al., 2002, Takemura et al., 2006). *mre11*Δ yeast cells are as sensitive to CPT as *rad52*Δ cells (**Figure 1**) and the combinations of *mre11*Δ or *rad52*Δ with *rad3-102* are lethal (Moriel-Carretero & Aguilera, 2010a), suggesting that Mre11 is absolutely required for HR-mediated replication restart. Thus, the results described in human HCT116 cells are reminiscent of the Mus81-dependent DSBs we have observed in *rad52*Δ cells, which likely represent the cleavage of blocked or reversed forks that could not undergo restart through HR (**Figure 7**) (Lemacon et al., 2017).

**Figure 7.**
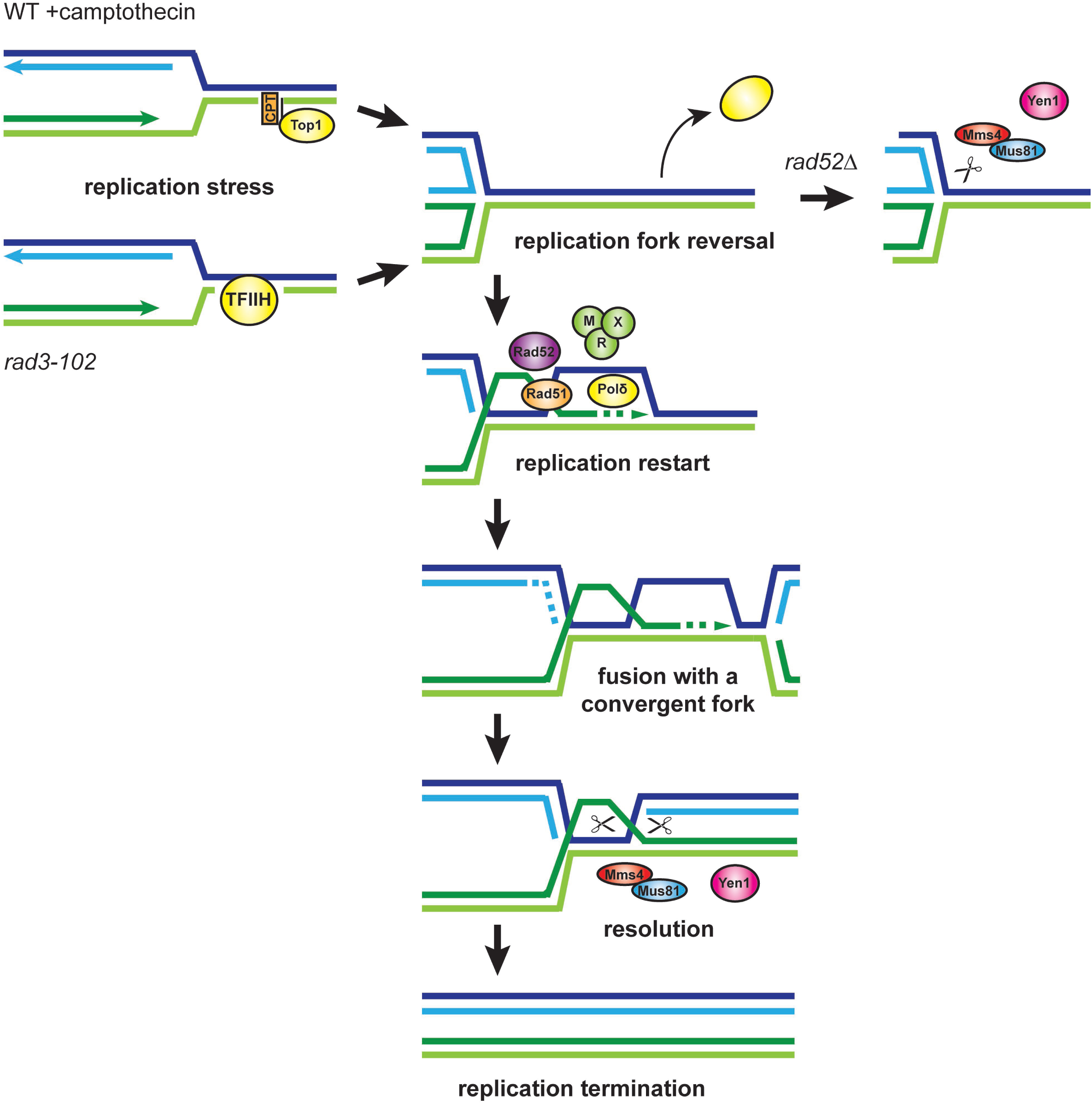
Proposed model for HR-mediated restart of replication. Replication fork block by CPT and in rad3-102 cells would result in fork reversal, favoring the removal of the replication impediment. The tip of the reversed fork would serve as a substrate for the invasion of the parental duplex by HR, requiring Rad52, Rad51, the MRX complex and the DNA polymerase δ (including Pol32). Replication could restart by BIR until the merging of the D-loop with a convergent fork, producing a Holliday junction. Resolution by Mus81 (or Yen1) would promote replication termination. Mus81 and Yen could also cleave replication forks that could not be restarted by HR.

Our results are consistent with a recent study on the survival to an inducible, replication-born, one-ended DSB, which primarily depends on Rad52 and Rad51 (Mayle et al., 2015). Since we did not detect DSBs in wild-type CPT-treated and in *rad3-102* cells, we favor the hypothesis that replication restart in our systems would use the tip of reversed forks as substrates for HR, as proposed in fission yeast and in mammalian cells (Teixeira-Silva et al., 2017, Zellweger et al., 2015). This would explain why the absence of the Ku complex, which is normally recruited to double-stranded DNA ends, could supress the resection defect caused by the lack of Mre11 nuclease activity in cells treated with CPT or containing the *rad3-102* allele **(Figure 1)** (Foster et al., 2011). Additionally, replication intermediates corresponding to fork reversal were observed by 2D-gel analyses in *rad3-102* cells (Moriel-Carretero & Aguilera, 2010a). The occurrence of fork reversal in *rad3-102* cells would suggest that fork reversal upon Top1 poisoning would not be the consequence of the accumulation of topological stress ahead of replication forks (Jakobsen et al., 2019a, Koster et al., 2007, Ray Chaudhuri et al., 2012). DNA end resection of reversed forks would involve the Mre11 and Exo1 nucleases (**Figure 1**) (Ait Saada et al., 2017, Kim et al., 2017, Tsang et al., 2014) and Rad52 and Rad51 would promote the efficient invasion of the parental duplex (**Figure 7**).

We had previously reported that replication restart in the absence of Rad51 in *rad3-102* cells can still be compensated by the presence of Pol32, a non-essential subunit of DNA polymerase δ (Moriel-Carretero & Aguilera, 2010a). The repair of replication-born DSBs by sister-chromatid recombination in a plasmid-based system confirmed the existence of a Rad51-independent Pol32-dependent pathway (Munoz-Galvan et al., 2013). A similar pathway, termed MIDAS, has been described in mammalian cells and depends on RAD52 and POLD3, the homolog of Pol32, but not on RAD51 (Bhowmick et al., 2016, Costantino et al., 2014, Sotiriou et al., 2016). Here, we have shown that Pol32 also compensates the absence of Rad51 for CPT resistance and DSB repair by BIR (**Figure 1**). These results have led us to propose that HR-mediated replication restart occurs by a BIR-like mechanism. The absence of Pol32 alone weakly affects cell survival in response to CPT and in *rad3-102* background (**Figure EV1A**) (Moriel-Carretero & Aguilera, 2010a, Vance & Wilson, 2002). Likewise, Pol32 is not primarily required for the repair of broken replication forks in the Flp-nick system (Mayle et al., 2015). The Pif1 DNA helicase is required for processive DNA synthesis during BIR (Saini et al., 2013, Wilson et al., 2013). However, the absence of nuclear Pif1 in *pif1-m2* mutants does not sensitize cells to CPT (Mayle et al., 2015), nor leads to lethality in *rad3-102 rad51*Δ cells (unpublished observations). These results suggest that processive BIR is not needed after the restart of blocked or broken forks by HR. Because DNA replication is started from various origins along chromosomes, blocked replication forks could be restarted by BIR but rapidly stopped by a convergent fork (**Figure 7**) (Mayle et al., 2015). The presence of backup replication origins in eukaryotic genomes, which are fired to ensure the completion of DNA replication, questions the benefit of fork restart. Fork restart could be essential in genomic regions poor in replication origins, such as common fragile sites (Debatisse & Rosselli, 2019). Additionally, processing of blocked forks by HR could protect nascent strands from degradation until encountering a convergent fork. Indeed, in the absence of Rad52 or BRCA2, nascent DNA strands at blocked or broken forks are extensively degraded by nucleolytic activities (Ait Saada et al., 2017, Jakobsen et al., 2019b, Lemacon et al., 2017, Schlacher et al., 2011), preventing the merging with a convergent fork and leading to mitotic catastrophes (Ait Saada et al., 2017).

Mus81 is required for cell survival in response to CPT and in the *rad3-102* background. (**Figure 2**). Our molecular analyses show that Mus81 is not required for HR-mediated restart of blocked forks but rather for the processing of joint molecules after the bulk of DNA synthesis in G_2_/M phase (**Figure 4** and **Figure 5**). After the restart of a broken fork by BIR, it has been proposed that Mus81 could cleave the migrating D-loop in order to limit the mutagenesis associated with BIR synthesis and re-establish a stable fork structure (Mayle et al., 2015). Our data argue against the possibility that Mus81 could fulfil this role during the S phase because Mus81 is not activated in S phase by the replication stress caused by CPT or *rad3-102* and the absence of Mus81 does neither affect nor ameliorate, the replication progression in these conditions. One possibility is that HR-mediated replication restart is initiated in S phase but the priming of DNA synthesis is delayed until G_2_/M, when Mus81 catalytic activity is enhanced by the hyper-phosphorylation of its regulatory partner Mms4. BIR initiated in G_2_/M also shows a delay between strand invasion and the initiation of DNA synthesis within the D-loop (Malkova et al., 2005). In the context of fork restart, this delay would give the opportunity to a convergent fork to reach the blocked forks “stabilized” by HR to ensure the completion of replication without challenging genome stability. The fusion of a D-loop with a convergent replication fork would lead to the formation of a nicked Holliday junction, for which the Mus81 SSE has a high affinity *in vitro* (Ehmsen & Heyer, 2009, Schwartz et al., 2012). Thus, we propose that the essential function of Mus81 after the restart of blocked forks is to process nicked Holliday junctions at termination sites (**Figure 7**).

In conclusion, our work describes how cells deal with blocked replication forks dispersed in the whole genome by Top1 poisoning. HR factors Rad52 and Rad51 orchestrate the restart of replication by a BIR-like mechanism and Mus81 promotes the termination of restarted forks with convergent forks. Because incomplete NER reactions in *rad3-102* cells are also occurring in wild type cells (Andersen et al., 2016) and Top1ccs have been found to accumulate naturally (Stingele et al., 2014), this mechanism is likely to be general in response to other replication blocks requiring HR to restart replication. Finally, the method we have described to use CPT in yeast cell cultures will allow performing a deeper characterization of Top1 poisoning by CPT at the molecular level. This will indubitably have important implications for understanding the effects of CPT as a chemotherapeutic agent.

## MATERIALS AND METHODS

### Yeast strains

All *Saccharomyces cerevisiae* yeast strains used in this study are in W303-1aR5 background (*his3-11, 15 leu2-3, 112, trp1-1 ura3-1 ade2-1 can1-100 RAD5*) unless indicated and are listed in table EV1. Deletion mutants were either obtained by the PCR-based gene replacement method (verified by PCR and drug sensitivity assays) or by genetic crosses (verified by tetrad analysis).

### Sensitivity assays to camptothecin (CPT) and ultra-violet irradiation (UV)

Cells from mid-log cultures were counted using a CASY^®^ (OLS system) and concentrated to 1×10^8^ cells/mL, 10µL of 10-fold serial dilutions were spotted on rich YPAD plates +/- CPT ((S)-(+)-Camptothecin, Sigma #C9911) and plates were incubated for 3 days at 30°C. To assay UV sensitivity, 10-fold serial dilutions were spotted on YPAD plates and plates were irradiated with UV-C in a Bio-Link™ BLX crosslinker and incubated for 3 days at 30°C in darkness. Three to four independent biological replicates have been performed for each sensitivity assay.

### Cell cycle progression analyses

Overnight mid-log cultures at 7×10^6^ cells/mL were synchronized in G_1_ with α-factor (0.5 µg/mL) in YPD medium for 2 to 3 hours at 30°C. For UV-induced DNA damage, G_1_-synchonized cells were resuspended in water onto Petri dishes as a 4mm-deep cell suspension, irradiated with UV-C in a Bio-Link™ BLX crosslinker, resuspended again in YPD medium with α-factor, incubated for 2 more hours in darkness and released into S phase by addition of pronase (50 µg/mL). For CPT-induced DNA damage, G_1_-synchonized cells were washed with MPD+SDS medium (0.17% yeast nitrogen base, 0.1% L-proline, 2% glucose and 0.003% SDS) (Pannunzio et al., 2004), resuspended in MPD+SDS medium supplemented with α-factor and DMSO or CPT, incubated for 1 more hour and released into S phase by addition of pronase (50 µg/mL). Samples (430µL) were taken every 10 or 15 minutes and fixed with 100% ethanol for subsequent flow cytometry analysis. Cells were centrifuged for 1 minute at 16.000 RCF, resuspended in 50 mM sodium citrate buffer containing 10 µl of RNase A (20 mg/ml, Qiagen 76254) and incubated for 2 hours at 50°C. Then, 10 µl of proteinase K (Sigma, P6556) were added for further incubation 2 hours at 50°C. Aggregates of cells were dissociated by sonication. 20 µl of cell suspension were incubated with 200 µl of 50 mM sodium citrate buffer containing 4 µg/mL propidium iodide (Sigma) for at least 2 hours in darkness. Data were acquired on a MACSQuant analyzer (Miltenyi Biotech) and analyzed with FlowJo software. Two to four independent biological replicates have been performed for each cell cycle progression analysis.

### DNA combing

DNA combing was performed essentially as described (Bianco et al., 2012). EdU incorporation and uptake in cells was realized in strains bearing 7 integrated copies of the *Herpes simplex* virus thymidine kinase (HSV-TK) and the human equilibrative nucleoside transporter 1 (hENT1) on a plasmid. Overnight mid-log cultures at 5×10^6^ cells/mL in YPD medium were washed with MPD +SDS medium and resuspended in MPD +SDS with DMSO or 50µM CPT, incubated for 2 hours and pulse-labeled with 50µM EdU for 20 minutes. DNA fibers were combed on silanized coverslips. Total DNA was detected with YOYO^®^-1 iodide (Molecular Probes Y3601) and EdU-labeled DNA was detected by Click chemistry using 20µL of the following mix per coverslip: 16.7 µL of H_2_O, 0.8 µL of 100 mM Copper sulfate, 0.5 µL of 0.5 mg/mL Alexa Fluor^®^ 555 Azide (Molecular Probes A20012) and 2 µL of 100 mM Sodium ascorbate. Coverslips were incubated for 1 hour at 60°C in a humid chamber in obscurity. Images were recorded on a Leica DM6000 microscope equipped with a CoolSNAP HQ CCD camera (Roper Scientific) and processed as described (Pasero et al., 2002). For statistical analysis, we used a Mann-Whitney test to compare, in each case, two unpaired groups with no Gaussian distribution. Two to three independent biological replicates were performed.

### Protein analyses

For time course experiments, proteins were extracted from cell pellets with acid-washed glass beads in a denaturing cracking buffer (8M Urea, 5% SDS, 40 mM Tris-HCl pH 6.8, 0.1 mM EDTA, 0.4 mg/mL bromophenol blue, 50 mM NaF, 150 mM β-mercaptoethanol, 2 mM PMSF) supplemented with cOmplete™ (Roche) protease inhibitors, 20 minutes at 70°C under permanent agitation at 1400 rpm in a Thermomixer^®^ (Eppendorf). Extracts were cleared by centrifugation, separated in Nupage^®^ 3-8% (Invitrogen) polyacrylamide gels and blotted on PVDF membranes using the Trans-Blot^®^ Turbo™ system (Bio-Rad). Two to three independent biological replicates have been performed for each time course experiment. For immunodetection, the following antibodies were used: anti-FLAG^®^ M2 (Sigma-Aldrich, F1804), anti-Clb2 (Santa Cruz Biotech, y-180) and anti-tubulin YOL1/34 (Abcam, ab6161).

### Pulsed-field gel electrophoresis

Agarose plugs containing chromosomal DNA were made as described (Tourrière et al., 2017). Chromosomes were separated at 13°C in a 0.9% agarose gel in TBE 0.5x using a Rotaphor apparatus (Biometra) using the following parameters: interval from 100 to 10 seconds (logarithmic), angle from 120 to 110° (linear), voltage 200 to 150 V (logarithmic). The gel was subsequently stained with ethidium bromide, and transferred to Hybond XL (GE Healthcare). Quantification of chromosome intensity was performed with Imaje J software after Southern blotting and hybridization using a radioactive probe specific for chromosome IV (*ARS453*) or VIII (*RRM3* gene), using a PhosphorImager (Typhoon Trio, GE). Two independent biological replicates were performed.

### Microscopy analyses

For the visualization of CPT inside cells, images were recorded on a Leica DM6000 microscope after excitation of CPT compound with a 350 nm UV light. Fluorescence quantification was performed with Image J software. For the analysis of spontaneous Rad52 foci, non-fixed nuclei from exponentially growing cell cultures bearing plasmid pWJ1344 were stained with DAPI and Rad52-YFP foci were counted in S-G_2_ cells. For the kinetic analyses of CPT-induced Rad52 foci, non-fixed nuclei were visualized thanks to mCHERRY-Pus1 and Rad52-YFP foci were counted in all cells. Three independent biological replicates were performed.

### 2D gel electrophoresis

For DNA extraction, 50 mL of the desired cultures were collected in Falcon tubes containing 500μL 10% Sodium Azide and kept in ice till processing. Cells were washed with 5 mL of chilled water and carefully resuspended in 1 mL of 1M sorbitol-10mM EDTA pH 8-0,1% β-mercaptoethanol-2 mg/mL Zymoliase 20T, and then incubated at 30°C during one hour under soft agitation. After centrifugation, the sferoplast pellet was washed with 500 μL of cold water and then broken with 400 μL of cold water plus 500 μL of 1.4M NaCl-100mM Tris-Cl pH 7.6-25mM EDTA pH 8-2% CTAB and incubated for thirty minutes at 50°C with 40 μL 10 mg/mL RNase. Next, 40 μL 20 mg/mL Proteinase K were added and incubation performed overnight under very soft agitation. After centrifugation, pellet and supernatant were treated separately. The supernatant was extracted with 500 μL (24:1) Chloroform:Isoamyl Alcohol and DNA precipitated with two volumes of 50 mM Tris-Cl pH 7.6-0 mM EDTA pH 8-1% CTAB and further resuspended in 250 μL 1.4M NaCl-1 mM EDTA pH 8-10 mM Tris-Cl pH 7.6. The original pellet was resuspended in 400 μL 1.4M NaCl-1 mM EDTA pH 8-10 mM Tris-Cl pH 7.6 and incubated during one hour at 50°C, extracted with 200 μL (24:1) Chloroform:Isoamyl Alcohol and rejoined with the supernatant DNA. The whole sample was precipitated then with one volume room temperature isopropanol, centrifuged, washed with 70% ethanol and resuspended in 100 μL 10mM Tris-Cl pH 8. DNA was digested using 200 U of each *Eco*RV and *Hind*III during 5 hours and NaCl-precipitated. First dimension electrophoresis was carried out at room temperature in 0.4% agarose gels at 40V during twenty hours in 1x TBE, next stained with ethidium bromide and a band comprised between 2 and 12 kb was cut and rotated 90° for the second dimension electrophoresis. Second dimension electrophoresis was carried out at 4°C in 1% agarose gels containing 0.34 μg/mL ethidium bromide at 130V during 12 hours in 1x TBE containing 0.34 μg/mL ethidium bromide. Gels were treated and transferred by standard procedures. For hybridization, coordinates of α^32-^P PCR probes were 37883-41883 for *ARS305* and 57903-61158 for region C on chromosome III. Signals were acquired using a Fujifilm FLA-5100 PhosphorImager. Two independent biological replicates were performed.

### BIR assay

Exponentially growing cells in YPR (2% Raffinose) medium were plated on rich medium containing 2% glucose (YPD) or 2% galactose (YPG) and incubated 3 days at 30°C. Colonies on YPG plates were replica-plated onto synthetic complete (SC) medium lacking lysine. BIR frequencies were determined by dividing the number of Lys+ by the number of YPD cfu. For statistical analysis, we used an unpaired Mann-Whitney t test. Three to six independent biological replicates were performed for each strain.

## ACKNOWLEDGEMENTS

We thank L.S. Symington, J.H. Petrini, S.C. West, W.D. Heyer, J.A. Tercero and D. Branzei for yeast strains and reagents. We thank Etienne Schwob and the DNA combing facility of Montpellier for providing silanized coverslips and the Montpellier RIO Imaging microscopy and flow-cytometry facility.

## FUNDINGS

This work was funded by grants from the “Agence Nationale pour la Recherche” (ANR), the “Institut National du Cancer” (INCa), and the “Ligue contre le Cancer” (équipe labellisée) to PP and from the Spanish Ministry of Economy and Competitiveness (BFU2013-42918P, and Consolider Ingenio 2010 CSD2007-015), the “Junta de Andalucia” (BIO-1238 and CVI-4567), and the European Union (FEDER) to AA. BP was supported by fellowships from the “Fondation Recherche Médicale” (SPF20121226243) and from the Spanish Ministry of Economy and Competitiveness (JCI 2009-04101). MMC was recipient of a predoctoral FPU training grant from the Spanish Ministry of Economy and Competitiveness.

## AUTHOR CONTRIBUTIONS

Conceived and designed the experiments: BP MMC AA PP. Performed the experiments: BP MMC TV. Analyzed the data: BP MMC AA PP. Wrote the paper: BP MMC AA PP.

## CONFLICT OF INTEREST

The authors declare that they have no conflict of interest.

## EXPANDED VIEW FIGURE LEGENDS

**Expanded View figure 1.** DNA repair is similar in CPT-treated and *rad3-102* cells.

**(A)** CPT sensitivity assayed by 10-fold serial dilutions of different mutant combinations on YPAD plates.

**(B) (C)** UV sensitivity assayed by 10-fold serial dilutions of different mutant combinations with *rad3-102* allele on YPAD plates after exposure to the indicated UV-C doses.

**Expanded View figure 2.** Yen1 backs up Mus81 in CPT-treated and *rad3-102* cells.

**(A)** CPT sensitivity assayed by 10-fold serial dilutions of different mutant combinations between *mus81*Δ and *yen1*Δ on YPAD plates.

**(B)** UV sensitivity assayed by 10-fold serial dilutions of *yen1*Δ in combination with *rad3-102* allele on YPAD plates after exposure to the indicated UV-C doses.

**(C)** Synthetic combinations of *rad3-102 mus81*Δ with *yen1*Δ and *pol32*Δ. Tetrads dissected on YPAD medium are shown. Triangles indicate lethality.

**Expanded View figure 3.** DNA repair is similar in CPT-treated and *rad3-102* cells.

**(A)** Visualization of CPT-containing yeast cells. Exponentially growing wild type cells in either MAD (minimal-ammonium-dextrose) or MPD +SDS (minimal-proline-dextrose + 0.003% SDS) medium were incubated with DMSO or 50 µM CPT for 30 min and visualized by fluorescence microscopy. Scale bar = 5 µm.

**(B)** Quantification of fluorescence emitted by CPT in cells grown as described in (A). A.U. arbitrary units.

**(C)** Analysis of DNA content by flow cytometry of G_1_ phase synchronized wild type cells and further released into S phase. Cells were synchronized in G_1_ with α –factor in either MAD or MPD +SDS medium, treated with DMSO or 50 µM CPT, let in G_1_ for 30 min, and released into S phase.

Asterisks indicate the progression of cells in S phase.

**Expanded View figure 4.** Mus81 is not required for replication fork progression in *rad3-102* cells.

Analysis of replicated DNA tracks length by single-molecule DNA combing in wild type, *mus81*Δ, *rad3-102* and *rad3-102 mus81*Δ cells. Exponentially growing cells pulse-labeled with 50 µM EdU for 20min. DNA fibers were combed on silanized coverslips and EdU-labeled DNA was detected by Click chemistry. Graph depicts the distribution of EdU tracks length in kb. Box and whiskers indicate 25-75 and 10-90 percentiles, respectively. Median EdU tracks length is indicated in kb. Asterisks indicate the *P*-value of the Man-Whitney unpaired t test, **** P-value <0.0001, ** P-value <0.01. Representative images of DNA fibers are shown. Red and white: EdU, green: DNA.

**Expanded View figure 5.** Mms4 phosphorylation in G_2_/M is required for recombination intermediates processing after replication restart.

**(A)** Analysis of Rad52 foci formation. Cells with Rad52-YFP foci were scored in exponentially growing wild-type *mus81*Δ, *rad3-102* and *rad3-102 mus81*Δ cells. Mean and SD of cells with Rad52 foci for three independent experiments are shown. An illustrative image is shown with Rad52 foci indicated by white arrows.

**(B)** Mms4-Flag10 phosphorylation analyzed by immunoblot in *rad52*Δ cells exposed to CPT. *rad52*Δ cells were synchronized in G_1_ with α-factor, treated with DMSO or 50 µM CPT, let in G_1_ for 1 h, and released into S phase. Cells were collected at the indicated time points and Mms4 was immunodetected with Flag antibodies. Clb2 immunodetection serves as a marker for G_2_ phase entry. Flow cytometry profiles corresponding the experimental setup are shown.

**(C)** Mms4-Flag10 phosphorylation analyzed by immunoblot in wild type and *rad3-102* cells. Cells were synchronized in G_1_ with α-factor, untreated or irradiated with 30 J/m^2^ UV-C, let in G_1_ for 2 h, and released into S phase. Cells were collected at the indicated time points and Mms4 was immunodetected with Flag antibodies. Clb2 immunodetection serves as a marker for G_2_ phase entry. Flow cytometry profiles corresponding the experimental setup are shown.

**(D)** CPT sensitivity assayed by 10-fold serial dilutions of wild type and *mms4* mutants.

**Expanded View figure 6.** DSBs are not detected in wild type CPT-treated and *rad3-102* cells.

**(A)** Analysis of DNA content by flow cytometry of G_1_ phase synchronized wild type and *mus81*Δ cells and further released into S phase. Cells were synchronized in G_1_ with α-factor, treated with DMSO or 100 µM CPT, let in G_1_ for 1 h, and released into S phase. Asterisks indicate when cells start the following cell cycle.

**(B)** Pulsed-field gel electrophoresis (PFGE) analysis of wild type cells in response to CPT. Wild type and *mus81*Δ cells were synchronized in G_1_ with α-factor, treated with DMSO or 200 µM CPT, let in G_1_ for 1 h, and released into S phase. Cells were collected at the indicated time points. DNA contents was analyzed by flow cytometry and the DNA extracted in agarose plugs was analyzed by PFGE. The agarose gel has been stained with ethidium bromide. zeo, wild type cells released from the G1 arrest in the presence of 1 mg/mL Zeocin for 1 h. Zeocin-induced DSBs are indicated by a vertical bar.

**(C)** PFGE analysis of wild type and *rad3-102* cells. Wild type and *rad3-102* cells were synchronized in G_1_ with α-factor, untreated or irradiated with 20 J/m^2^ UV-C, let in G_1_ for 2h, and released into S phase. Cells were collected at the indicated time points. DNA contents was analyzed by flow cytometry and the DNA extracted in agarose plugs was analyzed by PFGE. Upper panel: agarose gel stained with ethidium bromide; lower panels: Southern blot using a chromosome VIII specific probe. JMs, joint molecules accumulated in the gel wells. Quantification of JMs relative to the total amount of DNA is indicated for each time point.

## EXPANDED VIEW TABLE

**Expanded View table 1.** Yeast strains used in this study.

|  |  |
| --- | --- |
| YBP641 | *MATa bar1Δ ura3::URA3::GPD-TK<sub>7</sub>* |
| YPB642 | *MATa bar1ΔNatMX6 rad3-102::HphMX6 ura3::URA3::GPD-TK<sub>7</sub>* |
| YBP643 | *MATa bar1ΔNatMX6 mus81ΔKanMX4 ura3::URA3::GPD-TK<sub>7</sub>* |
| YBP645 | *MATa bar1Δ rad3-102::HphMX6 mus81ΔKanMX4 ura3::URA3::GPD-TK<sub>7</sub>* |
| YBP782 | *MATa bar1Δ mus81-dd ura3::URA3::GPD-TK<sub>7</sub>* |
| YBP785 | *MATa bar1Δ rad3-102::HphMX6 mus81-dd ura3::URA3::GPD-TK<sub>7</sub>* |
| YREC102-51 | *MATa rad3-102::HphMX6 mus81ΔKanMX4 rad51ΔKanMX4* |
| YBP1068 | *MATa bar1Δ rad3-102::HphMX6 rad51Δ::LEU2 ura3::URA3::GPD-TK<sub>7</sub>* |
| YBP1038 | *MATa RAD52-YFP RFA1-CFP* |
| YBP1043 | *MATa RAD52-YFP RFA1-CFP mus81Δ::NatMX6* |
| YBP699 | *MATa bar1Δ rad52Δ::klTRP1MX6 Mms4-His6-Flag10::KanMX4 ura3::URA3::GPD-TK<sub>7</sub>* |
| WS-M11 | *MATa mre11Δ::KanMX4* |
| WS-R50 | *MATa rad50ΔKanMX4* |
| WS-X2 | *MATa xrs2ΔKanMX4* |
| YBP665 | *MATa bar1Δ rad51Δ::HphMX6 ura3::URA3::GPD-TK<sub>7</sub>* |
| YBP667 | *MATa bar1Δ pol32Δ::NatMX6 ura3::URA3::GPD-TK<sub>7</sub>* |
| YBP669 | *MATa bar1Δ rad51Δ::HphMX6 pol32Δ::NatMX6 ura3::URA3::GPD-TK<sub>7</sub>* |
| YREC102-49 | *MATa mus81ΔKanMX4 rad51ΔKanMX4* |
| YREC102-47 | *MATa mus81ΔKanMX4 pol32ΔHphMX6* |
| YBP716 | *MATa bar1Δ mms4Δ::klTRP1MX6 ura3::URA3::GPD-TK<sub>7</sub>* |
| YBP759 | *MATa bar1Δ rad3-102::HphMX6 mms4Δ::klTRP1MX6 ura3::URA3::GPD-TK<sub>7</sub>* |
| YBP727 | *MATa bar1Δ pADH1-HA3-mms4-9A::HIS3* |
| YBP732 | *MATa bar1Δ mms4-14A-HA3::URA3* |
| YBP671 | *MATa bar1Δ Mms4-His6-Flag10::KanMX4 ura3::URA3::GPD-TK<sub>7</sub>* |
| YBP673 | *MATa bar1Δ rad3-102::HphMX6 Mms4-His6-Flag10::KanMX4 ura3::URA3::GPD-TK<sub>7</sub>* |
| YBP697 | *MATa bar1Δ yen1Δ::klTRP1MX6 Mms4-His6-Flag10::KanMX4 ura3::URA3::GPD-TK<sub>7</sub>* |
| YBP712 | *MATa bar1Δ::NatMX6 mus81Δ::KanMX4 yen1Δ::klTRP1MX6 ura3::URA3::GPD-TK<sub>7</sub>* |
| YREC105-24 | *MATa rad3-102::HphMX6 yen1Δ::KanMX4* |
| YBP1178 | *MATa bar1Δ mre11-3::URA3* |
| YBP1179 | *MATa bar1Δ yku70Δ::KanMX6* |
| YBP1180 | *MATa bar1Δ mre11-3::URA3 yku70Δ::KanMX6* |
| YBP1181 | *MATa bar1Δ rad3-102::HphMX6 mre11-3::URA3* |
| YBP1183 | *MATa bar1Δ rad3-102::HphMX6 yku70Δ::KanMX6* |
| YBP1185 | *MATa bar1Δ rad3-102::HphMX6 mre11-3::URA3 yku70Δ::KanMX6* |
| YBP1187 | *MATa bar1Δ exo1Δ::HIS3* |
| YBP1189 | *MATa bar1Δ mre11-3::URA3 exo1Δ::HIS3* |
| YBP1191 | *MATa bar1Δ exo1Δ::HIS3 yku70Δ::KanMX6* |
| YBP1193 | *MATa mre11-3::URA3 exo1Δ::HIS3 yku70Δ::KanMX6* |
| YBP1195 | *MATa bar1Δ rad3-102::HphMX6 exo1Δ::HIS3* |
| YBP1197 | *MATa rad3-102::HphMX6 mre11-3::URA3 exo1Δ::HIS3* |
| YBP1199 | *MATa bar1Δ rad3-102::HphMX6 exo1Δ::HIS3 yku70Δ::KanMX6* |
| YBP1201 | *MATa rad3-102::HphMX6 mre11-3::URA3 exo1Δ::HIS3 yku70Δ::KanMX6* |
| LSY2743-1.1 | *MATa-inc bar1Δ::LEU2 ade3::pGAL-HO ura3::URA3::GPD-TK<sub>7</sub> lys2Δ::NatMX4 avt2::lys-HOcs::KanMX6 cos9::TRP1-ys2* |
| LSY2753-3.8 | *MATa-inc bar1Δ::LEU2 ade3::pGAL-HO ura3::URA3::GPD-TK<sub>7</sub> lys2Δ::NatMX4 avt2::lys-HOcs::KanMX6 cos9::TRP1-ys2 pol32Δ::KanMX6* |
| LSY2824-21D | *MATa-inc bar1Δ::LEU2 ade3::pGAL-HO ura3::URA3::GPD-TK<sub>7</sub> lys2Δ::NatMX4 avt2::lys-HOcs::KanMX6 cos9::TRP1-ys2 rad51Δ::LEU2* |
| PP3475 | *MATa-inc bar1Δ::LEU2 ade3::pGAL-HO ura3::URA3::GPD-TK<sub>7</sub> lys2Δ::NatMX4 avt2::lys-HOcs::KanMX6 cos9::TRP1-ys2 pol32Δ::KanMX6 rad51Δ::HphMX6* |
| PP3476 | *MATa-inc bar1Δ::LEU2 ade3::pGAL-HO ura3::URA3::GPD-TK<sub>7</sub> lys2Δ::NatMX4 avt2::lys-HOcs::KanMX6 cos9::TRP1-ys2* |
|  | *pol32Δ::KanMX6 rad51Δ::HphMX6* |
| PP708 | *MATα ura3::URA3::GPD-TK<sub>7</sub> rev3Δ ::NatMX4* |
| YBP1203 | *MATa bar1Δ rad3-102::HphMX6 wss1Δ::KanMX4* |
| YBP1204 | *MATa bar1Δ rad3-102::HphMX6 wss1Δ::KanMX4* |
| YBP1206 | *MATa bar1Δ wss1Δ::KanMX4* |
| YBP1207 | *MATa tdp1Δ::NatMX6* |
| YBP1209 | *MATa bar1Δ rad3-102::HphMX6 tdp1Δ::NatMX6* |
| YBP1097 | *MATa bar1Δ rad3-102::HphMX6 NatMX6::G2-MUS81* |
| YBP1101 | *MATa NatMX6::G2-MUS81* |
| YBP816 | *MATa bar1Δ rad52Δ::klLEU2 mus81Δ::KanMX4* |

